# Spatial scale influences taxon conservation in the eukaryotic virome of a mosquito

**DOI:** 10.1101/2022.11.23.517782

**Authors:** Patricia Gil, Antoni Exbrayat, Etienne Loire, Ignace Rakotoarivony, Florian Charriat, Thierry Baldet, Michel Boisseau, Albane Marie, Benoît Frances, Gregory L’Ambert, Mohamed Bessat, Yehia Otify, Maria Goffredo, Giuseppe Mancini, Núria Busquets-Martí, Lotty Birnberg, Sandra Talavera, Carlos Aranda, Emna Ayari, Selma Mejri, Soufien Sghaier, Amal Bennouna, Hicham El Rhaffouli, Thomas Balenghien, Ghita Chlyeh, Ouafaa Fassi Fihri, Julie Reveillaud, Yannick Simonin, Marc Eloit, Serafin Gutierrez

## Abstract

Our knowledge on the diversity of eukaryotic viruses has recently undergone a massive expansion. This diversity could influence host physiology through yet unknown phenomena of potential interest to the fields of health and food production. However, the assembly processes in eukaryotic viromes of terrestrial animals remain elusive. This situation hinders hypothesis-driven tests of virome influence on host physiology. Here, we explore virome assembly at different spatial scales in the eukaryotic virome of the mosquito *Culex pipiens*. This mosquito is a vector of human pathogens worldwide. Several new aspects of virome assembly were unveiled through a sampling involving five countries in Africa and Europe and large sample sizes. A group of viruses was conserved in *C. pipiens* populations in all countries. This core group represented a relatively large and diverse fraction of the virome. However, core viruses were not shared by all host individuals in a given country, and their infection rates fluctuated between countries and years. Moreover, the distribution of co-infections in individual mosquitoes suggested random co-occurrence of certain core viruses. We also observed differences in the virome depending on geography, with viromes tending to cluster depending on the continent. Thus, our results unveil that taxon conservation in a eukaryotic virome changes with spatial scale. Thus, predictions on virome assembly seem possible at a large geographical scale in *C. pipiens*.

**IMPORTANCE:** The study of the eukaryotic virome of mosquitoes is an emerging research field. Beyond its fundamental interest, this field could lead to the development of control tools against the transmission of mosquito-borne human pathogens. However, we yet know little on the assembly patterns in the eukaryotic viromes of mosquitoes, as well as of terrestrial animals in general. This situation hampers the design of hypothesis-driven studies on the influence of the virome on pathogen transmission. Here, we have analyzed virome assembly in the mosquito vector *Culex pipiens* within and between countries in Africa and Europe. Our results show that integrating contrasted spatial scales allows to identify deterministic patterns in virome assembly. Such patterns can guide future studies of virome influence on mosquito physiology.

## Introduction

An impressive expansion in the known diversity of viruses has recently taken place thanks to metagenomics (1). This expansion has been followed by major improvements in the emerging field of community ecology of viruses. For example, viral communities or viromes have been shown to play a major role in the regulation of the carbon cycle in the oceans (2), or in bacteriome dynamics (3, 4). However, current knowledge is largely biased towards viromes of unicellular hosts. Our understanding of the ecology of the eukaryotic virome in terrestrial animals largely lags behind (5, 6). For example, we still know little about the potential deterministic patterns behind virome assembly in those organisms (6-10). This situation severely hampers, among others, studies on the influence of the eukaryotic virome on the physiology of those hosts.

The eukaryotic virome of mosquitoes has received considerable attention due to its potential influence on the transmission of human pathogens with a huge impact on health worldwide (11). As for most eukaryotic viromes, most studies on mosquito viromes have focused on describing taxonomic diversity (*e.g*., (9, 12-15)). Those studies have discovered an impressive number of new viruses. These viruses are mostly RNA viruses, the dominant group in terrestrial eukaryotes (16), and are thought to be mosquito commensals that do not infect vertebrates (17). Moreover, they are often supposed to be vertically transmitted, although other transmission modes have been observed (18-21). Interestingly, a few mosquito-specific viruses have been shown to influence the infection of mosquitoes by human pathogens under laboratory conditions (21-26).

The improvement in the known diversity of mosquito viruses contrasts with the paucity of studies on virome assembly patterns. For example, little is known on the potential influence of spatial scales on virome assembly (27, 28), a main question in community ecology (29, 30). The study of virome assembly across scales allows to infer whether specific viruses are associated to a host no matter the spatial scale, that is, whether a core virome exists. The identification of core taxa is often a main initial step in the study of microbial communities since core members are likely to have a key influence on host physiology (31, 32). To date, a fistful of studies has explored the potential existence of a core virome in different mosquitoes (9, 10, 33-39). Overall, certain viruses have been found repeatedly associated to a given mosquito species, suggesting that a cove virome may take place. However, current studies analyze a single spatial scale (9, 10, 33-35, 38, 39), or different but limited geographical ranges of the mosquito host (28). Moreover, sample sizes in those studies are usually low despite the usual large population sizes of mosquitoes (10, 34-36). This situation hampers to robustly elucidate whether a core virome is conserved across spatial scales.

Here, we analyze the influence of spatial scale on the eukaryotic virome of the common house mosquito *Culex pipiens* Linnaeus, 1758. This mosquito is the main vector of important human pathogens, like West Nile virus. Moreover, *C. pipiens* has nearly a worldwide distribution, probably through introductions from Europe and Africa (40). We have analyzed the virome of *C. pipiens* both between and within countries in Europe and Africa (Fig. 1). Our mosquito collection allowed to analyze between hundreds and thousands of individuals per country, and from the main habitats of *C. pipiens* in the study area. Using a blend of meta-transcriptomics (9, 12, 41) and PCR analyses, we revealed that taxon conservation decreased with spatial scale. That is, a group of viruses was shared between countries. However, a limited taxon conservation in the virome took place between mosquitoes within a country, along with random co-infection patterns. Moreover, differences in virome assembly were observed between European and African countries. Geographic structuration did not seem to originate by strict isolation of virus populations but rather from differences in relative abundancy within viromes.

**Figure 1.**
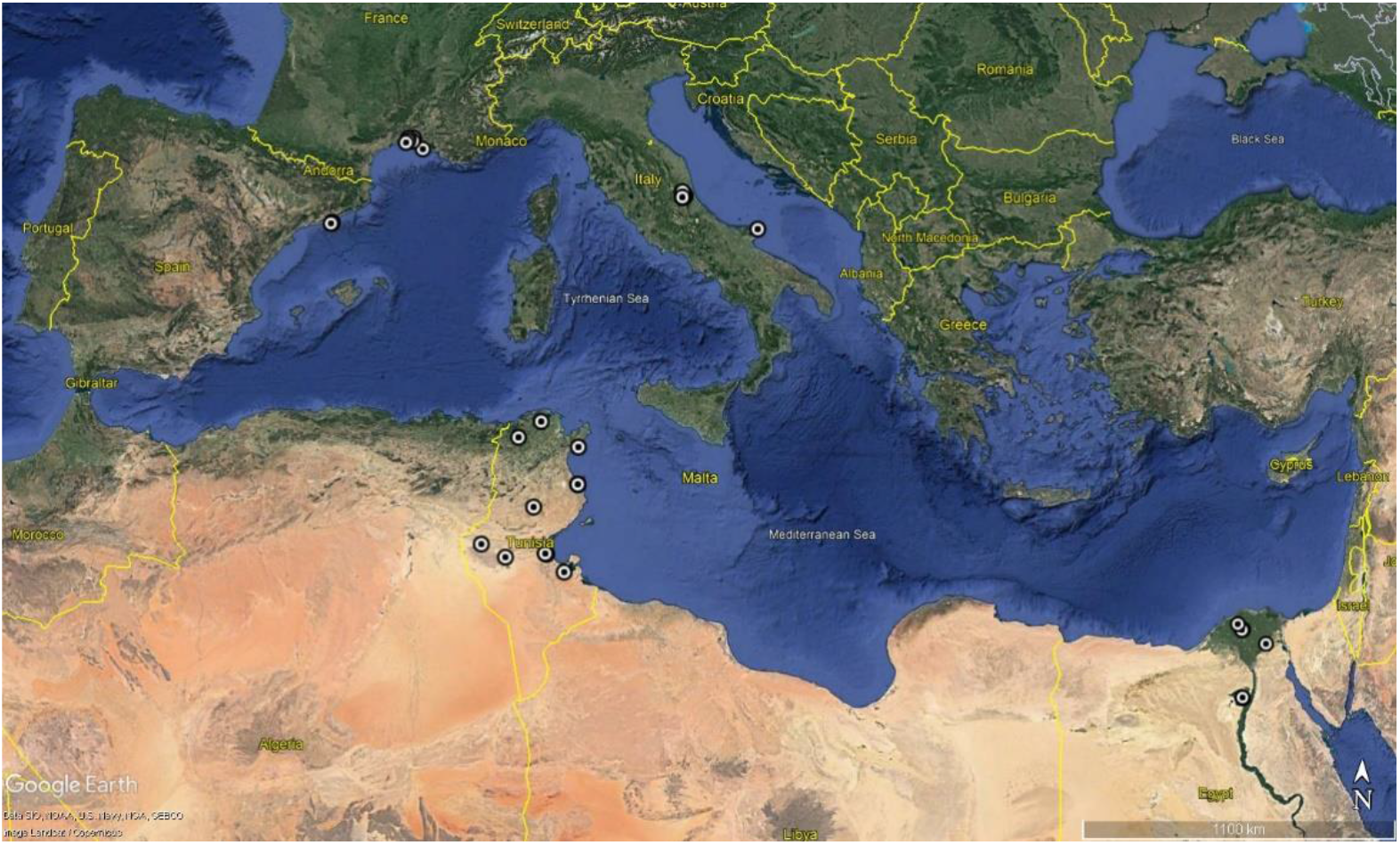
Overview of mosquito sampling sites in the Mediterranean Basin. Detailed information on the sites can be found in Fig. S7 and Table S4.

## Results

### High taxon conservation around the Mediterranean Basin

We used meta-transcriptomics (12) to characterize the eukaryotic virome of *C. pipiens* in five countries in Africa and Europe (Fig. 1). Two samples were generated per country, each comprising 711 females on average (Table S1). Samples were derived from different habitats to maximize the exploration of virus diversity (Table S1). Meta-transcriptomics analysis of the ten samples provided 715 million reads (Table S1). Among them, 41 million reads (5.7%) mapped to virus-like sequences associated with 324 viral operational taxonomic units (vOTUs) (36). The number of virus-like reads significantly differed between samples (Table S1; Wilcoxon signed rank exact test, *p* = 0.001). We did not detect a significant effect of raw reads or mosquito number on the yield of virus-like reads (Table S1; Spearman’s rank correlation, *p* > 0.076). Rarefaction curves showed that sampling effort, in terms of viral reads, was overall adequate (Fig. S1).

After quality filtering, the dataset comprised 201 vOTUs, distributed over 44 clusters (an arbitrary taxon similar to a Family, see Material and Methods), and 99.98% of the virus-like reads. The dataset was dominated by vOTUs related to viruses detected in arthropods (167 vOTUs, 99.90% of the virus-like reads). We were only interested in vOTUs potentially infecting *C. pipiens*. We thus discarded the vOTUs related to non-arthropod hosts. None of the non-arthropod vOTUs was found in more than six samples.

The final dataset consisted of 167 vOTUs and 36 clusters, mainly RNA viruses. A large proportion of the vOTUs had identities at the amino-acid level below 80% with their best hits in GenBank (Fig. S2). This situation suggests the presence of numerous new viruses, as usually observed in studies on the mosquito virome (10, 27, 28). Selected sequences were submitted to GenBank (BioProject PRJNA806751). Sequence submission was done with the Gsub tool, a new tool allowing easy annotation and submission of virus contigs (Supplementary Annex 1). Most names of new virus species were provided by children participating to the 2021 Science Days in Montpellier (France). A detailed description of this diversity is beyond the scope of this study and was not further explored here.

Cluster conservation was relatively large between libraries and countries (Fig. 2A). Ten clusters were present in all the libraries (27% of the clusters; hereafter referred to as “highly-conserved clusters”; Fig. 2A). Moreover, almost half of the clusters were present in all countries (18 clusters, Fig. 2A). Highly-conserved clusters tended to rank higher in terms of read abundance but some of them had reproducibly low frequencies in most libraries (*i*.e. < 0.1 %; *e.g*., see clusters Mononega_unclass and Picorna_unclass in Fig. 2B). More generally, read distribution among clusters was rather heterogeneous, with a few clusters providing most of the reads both globally and in each library (Fig. 2B).

**Figure 2.**
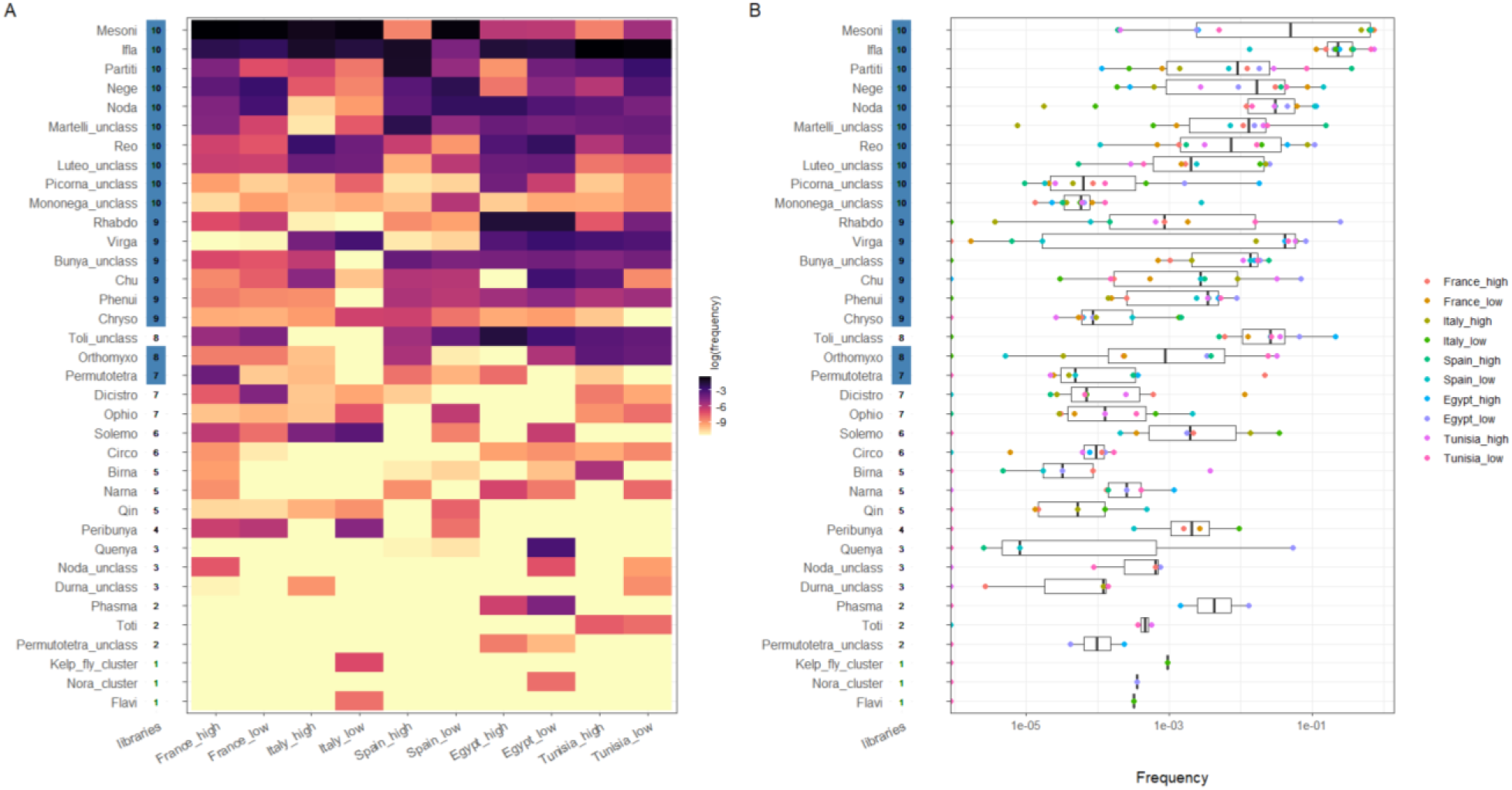
Structure of the eukaryotic virome of *Culex pipiens* in the Mediterranean Basin. (A) Heatmap showing the relative abundance of clusters among libraries. Clusters are ranked on the y axis first by number of libraries with presence and, then, by total read number. The number of libraries in which a given cluster was found is shown on the right of the cluster name. This number is underlined in blue if the cluster was detected in all countries. The darker the tile color, the closer the relative frequency of a given cluster to one. Libraries are shown on the x axis with the European countries on the right and the African ones on the left to facilitate comparison. The terms “high” and “low” on the library names stand for levels of habitat anthropization. (B) Boxplots with relative abundance of clusters. Clusters are ranked as in the A panel. Dots indicate frequency in a given library and the dot color stands for the library as shown in the legend.

We also observed a relatively high taxon conservation at the vOTU level (Fig. 3, Table S2). Seven out of the ten highly-conserved clusters included a vOTU that was present in all libraries (Fig. 3, Table S2). The three remaining clusters (Partiti, Mononega_unclass and Nege) had a vOTU present in at least eight libraries (Fig. 3, Table S2).

**Figure 3.**
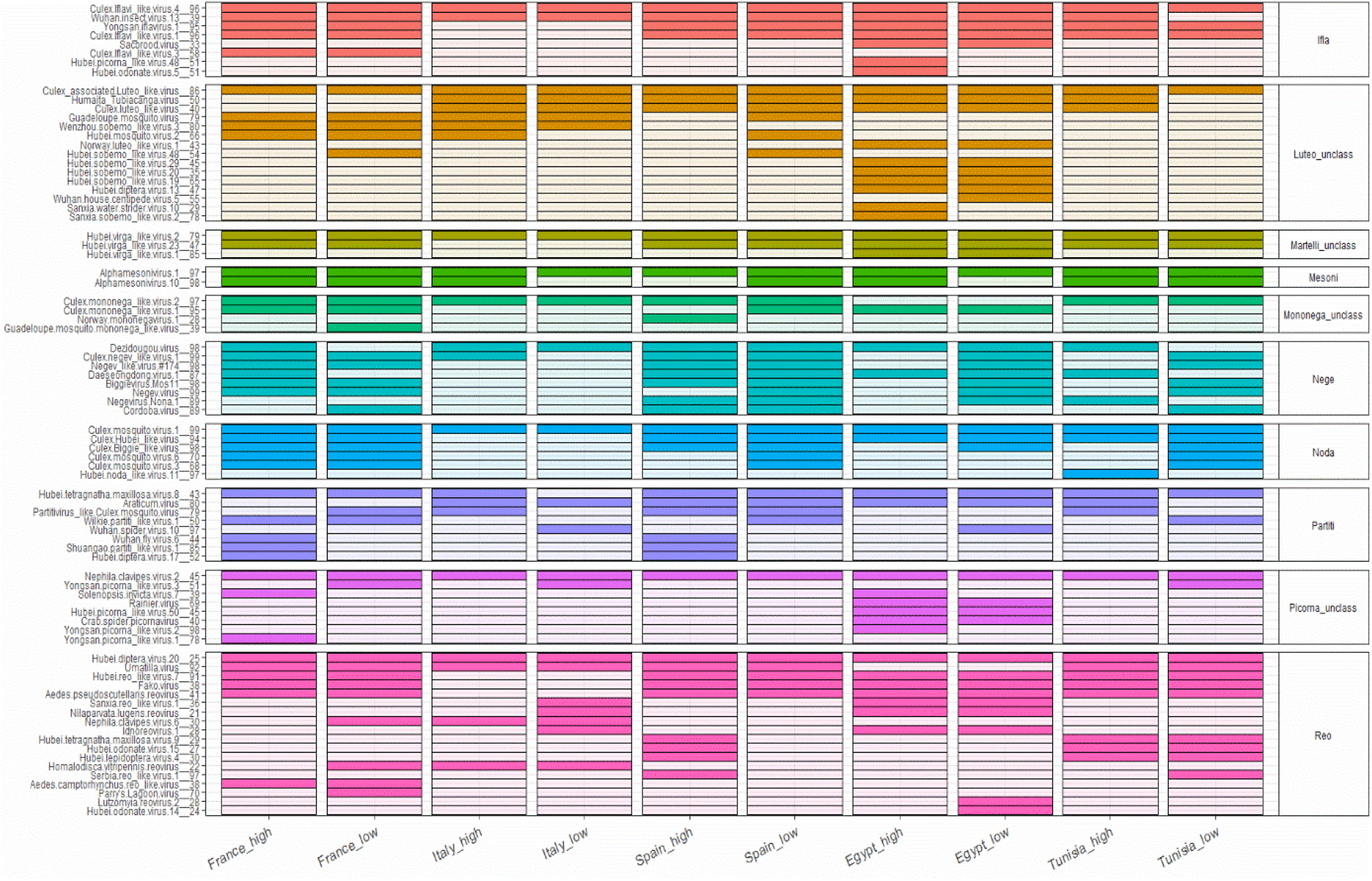
Prevalence of viral operational taxonomic units (vOTUs) from highly-prevalent clusters in the eukaryotic virome of *Culex pipiens* among libraries. The names of vOTUs and the associated clusters appear on the right and left of the heatmap, respectively. Colored tiles indicate detection in a given library. Tile color stands for cluster to facilitate visualization.

### Highly-conserved taxa are found associated to *Culex pipiens* in other regions

We analyzed if the highly-conserved taxa had been found associated to *C. pipiens* in other countries. Our dataset was compared to four studies from China, Morocco, Serbia and Sweden (10, 38, 48, 49). Despite important differences in methodology between studies (Table S3), we observed a relatively good conservation of highly-conserved clusters (Fig. 4). All clusters were detected in Morocco and Sweden. The study from China detected seven out of ten clusters despite of the fact that it only provides vOTUs that shared more than a 90 % identity at the amino-acid level with their best hit. Finally, the study from Serbia detected six clusters despite providing only those vOTUs for which complete RdRp genes or genomes had been obtained. Taxon conservation was also observed at the vOTU level. All highly-conserved clusters, except for the Partiti cluster, had vOTUs shared in other countries (Fig. 4).

**Figure 4.**
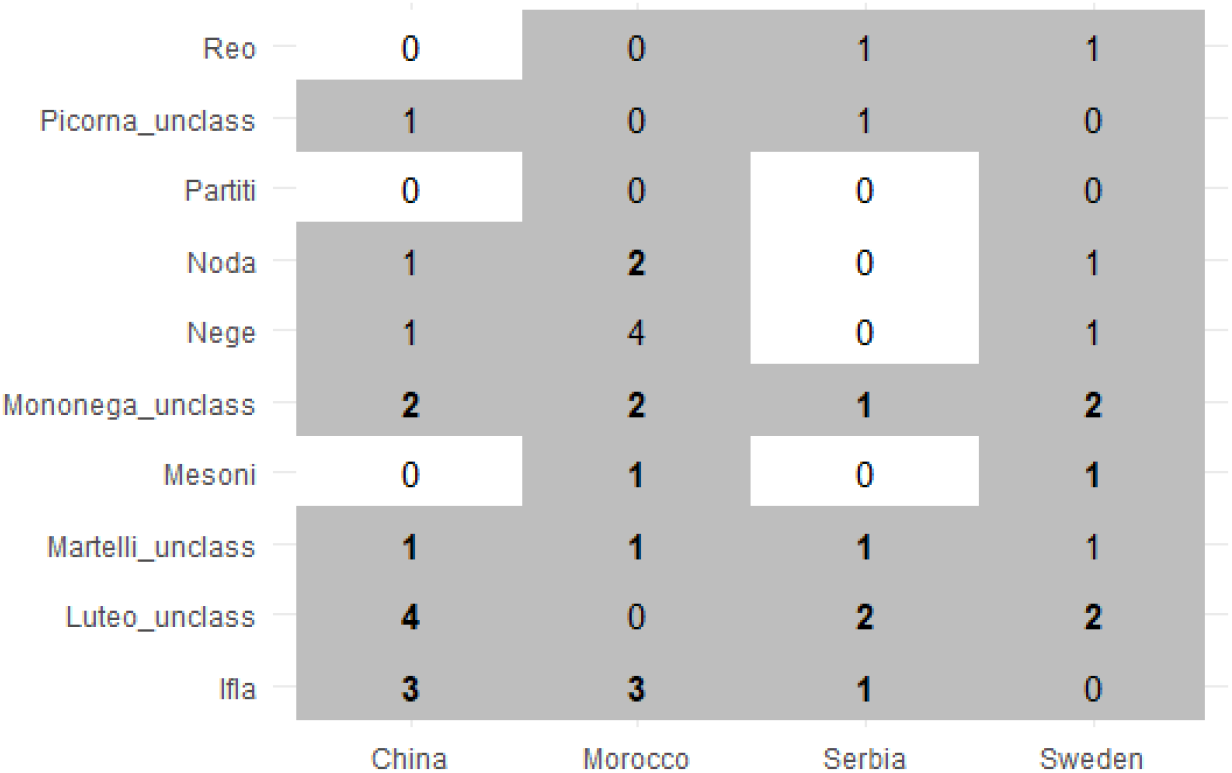
Taxon conservation in the virome of *Culex pipiens* between our study and studies from other countries. Grey tiles indicate highly-conserved clusters shared in our dataset and in a specific country (either China, Morocco, Serbia or Sweden). Numbers within tiles indicate the number of viral operational taxonomic units (vOTUs) of a given cluster also detected in our dataset. The vOTU number appears in bold if the vOTU with the highest prevalence in our dataset was also found in a given country (see Fig. 3 for those vOTUs).

### Taxon conservation between countries does not imply conservation between individual mosquitoes in a country

We explored whether taxon conservation also took place within each country. We estimated the infection rates of five vOTUs (Fig. S3), including two highly-conserved vOTUs, in each country. None of the vOTUs reached an infection rate above 10% in any of the countries (Fig. 5A). We observed significant differences in infection rates between vOTUs (Kruskal-Wallis, *p* = 0.014). More precisely, the infection rates of the highly-conserved Culex.Iflavi_like.virus.4__96 were significantly higher than those of Culex.Bunyavirus.2__98 and Culex.mosquito.virus.4__97 (Conover-Iman test, *p* = 0.010 and 0.002, respectively). Moreover, significant differences in infection rates were observed between countries for a given virus (Fig. 5A). We did not detect a significant effect of the country on the infection rates of all viruses (Kruskal-Wallis, *p* = 0.37).

**Figure 5.**
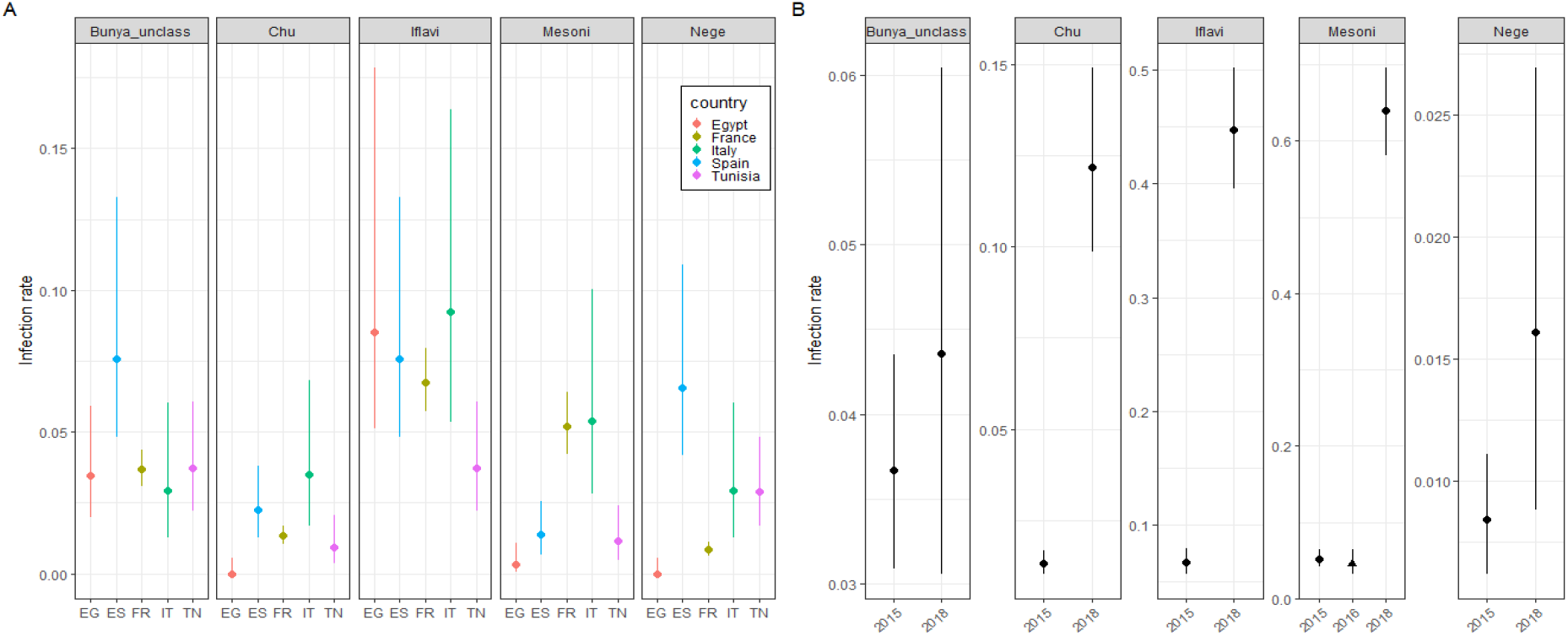
Infection rates of five viral operational taxonomic units (vOTUs) in *Culex pipiens* females. The five vOTUs were Culex.Bunyavirus.2__98 (Bunya_unclass), Culex.mosquito.virus.4__97 (Chu), Culex.Iflavi_like.virus.4__96 (Iflavi), Alphamesonivirus.1__97 (Mesoni) and Culex.negev_like.virus.1__99 (Nege). Infection rates are grouped in panels depending on the cluster of the vOTU. Dots indicate the maximum likelihood estimates of the infection rates (infected females per 100 females) and vertical segments stand for the 95% confidence intervals. (A) Infection rates of the five vOTUs in the five countries sampled in 2015. EG: Egypt, ES: Spain, FR: France, IT: Italy, TN: Tunisia. (B) Infection rates of the five virus taxa in France in 2015 and 2018. The infection rate of the vOTU in the Mesoni cluster in Morocco in 2016 is also included to facilitate comparison (marked with a triangle).

We then explored whether infection rates could change among years. To do so, 748 females collected in France in 2018 were screened for the five vOTUs. Infection rates in 2018 were often significantly higher than in 2015 (Fig. 5B). For example, Alphamesonivirus.1__97 and Culex.Iflavi_like.virus.4__96 showed infection rates close or above 50% in 2018. To further validate our results, we estimated the infection rate of Alphamesonivirus.1__97 in over 1 300 adult females of *C. pipiens* collected in another country and year (Morocco, 2016) (48). The infection rate lied below 10% in the Moroccan samples, like in the other five countries in 2015 (Fig. 5B). Overall, our results showed that the infection rates of highly-conserved vOTUs could be relatively low, and differed between countries and years.

### Frequent random co-infection in individual mosquitoes

We explored the potential non-random co-occurrence of different viruses within individual mosquitoes. We thus screened 184 individual females for the five vOTUs. Those females had been collected in France in 2018, the year with the highest infection rates for most vOTUs (Fig. 5). This analysis showed that 54% of the females were infected by at least two different vOTUs (Fig. 6A). Different combinations of vOTUs were observed among individuals (Fig. 6B). A probabilistic approach was used to infer co-occurrence patterns in vOTU pairs (58). Of the ten possible pairs, one pair was not included in the analysis because its expected co-occurrence was below 1% (the pair Culex.negev.like_virus.1__99/Culex.mosquito.virus.1__97). Only one pair out of the nine pairs tested showed an association deviating from random (positive association between Culex.Iflavi_like.virus.4__96 and Culex.Bunyavirus.2__96, *p* = 0.004; other pairs: *p* > 0.061).

**Figure 6.**
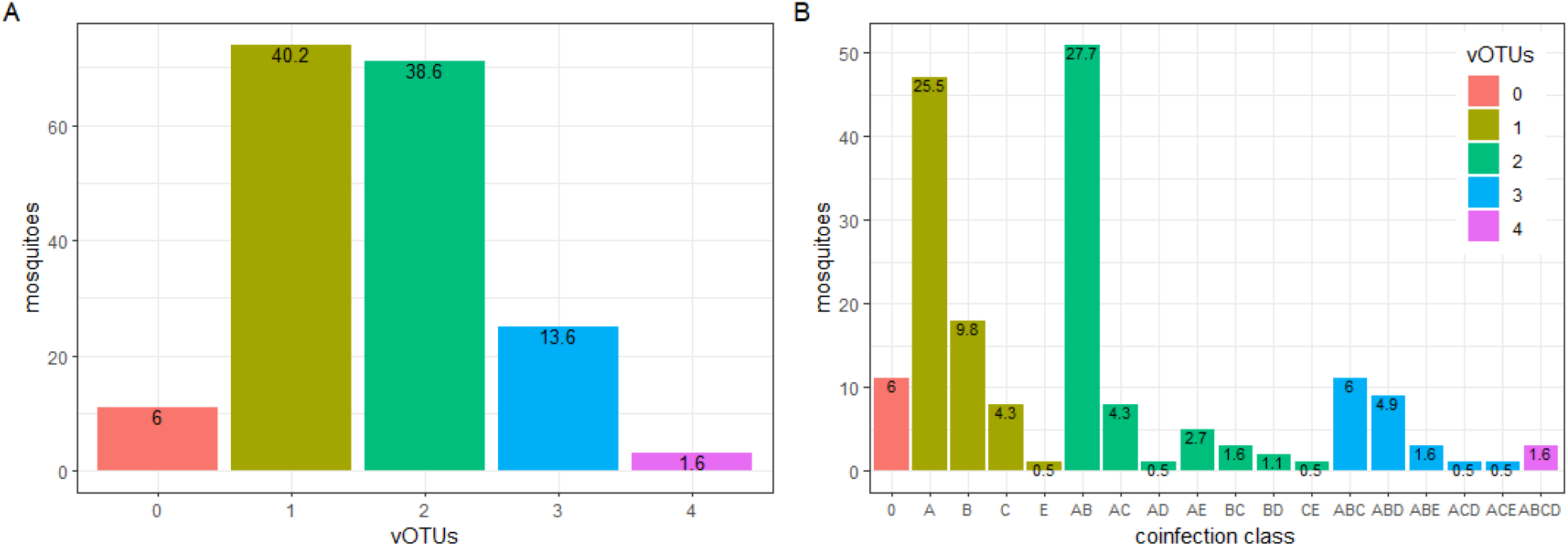
Distribution of viral operational taxonomic units (vOTUs) in individual *Culex pipiens* females. The five vOTUs were Culex.Bunyavirus.2__98 (Bunya_unclass), Culex.mosquito.virus.4__97 (Chu), Culex.Iflavi_like.virus.4__96 (Iflavi), Alphamesonivirus.1__97 (Mesoni) and Culex.negev_like.virus.1__99 (Nege). (A) Distribution of the number of vOTUs in individual mosquitoes. Bar color stands for the number of vOTUs per mosquito. The relative frequencies of each coinfection group are shown on top of the bars (in percentages). (B) Distribution of combinations of specific vOTUs (*i.e*. coinfection classes) found in individual *C. pipiens* females. Relative frequencies of each coinfection class are shown on top of the bars (in percentages). Bars are colored following the number of vOTUs per mosquito, as in panel A. Each capital letter stands for a given vOTU as follows. A: Alphamesonivirus.1__97 (Mesoni), B: Culex.Iflavi_like.virus.4__96 (Ifla), C: Culex.Bunyavirus.2__98 (Bunya_unclass), D: Culex.negev.like_virus.1__99 (Nege), and E: Culex.mosquito.virus.4__97 (Chu).

### Geography influences virome assembly

We explored potential factors behind the virome heterogeneity observed between samples (Fig. 2A). The habitat anthropization, the continent, and the number of mosquitoes or sampling sites per library did not show a significant influence on taxon richness or the Simpson and Shannon indexes (Kruskall Wallis test, *p* > 0.33; Fig. S4). Jaccard dissimilarities tended not to reach 0.5 but Bray-Curtis dissimilarities were often higher than 0.5 (Fig. S5). These results suggest that the assemblies often shared taxa although with different relative frequencies.

Next, we compared virome structure between libraries using non-parametric multidimensional scaling (NMDS) and Bray-Curtis dissimilarities, as well as hierarchical clustering (Fig. 7). Viromes tended to aggregate according to continent and country (Fig. 7). Clustering based on continent but not on habitat anthropization was further confirmed with a PERMANOVA (continent: R2 = 0.445, *p* = 0.01; anthropization level: R2 = 0.062, *p* = 0.35).

**Figure 7.**
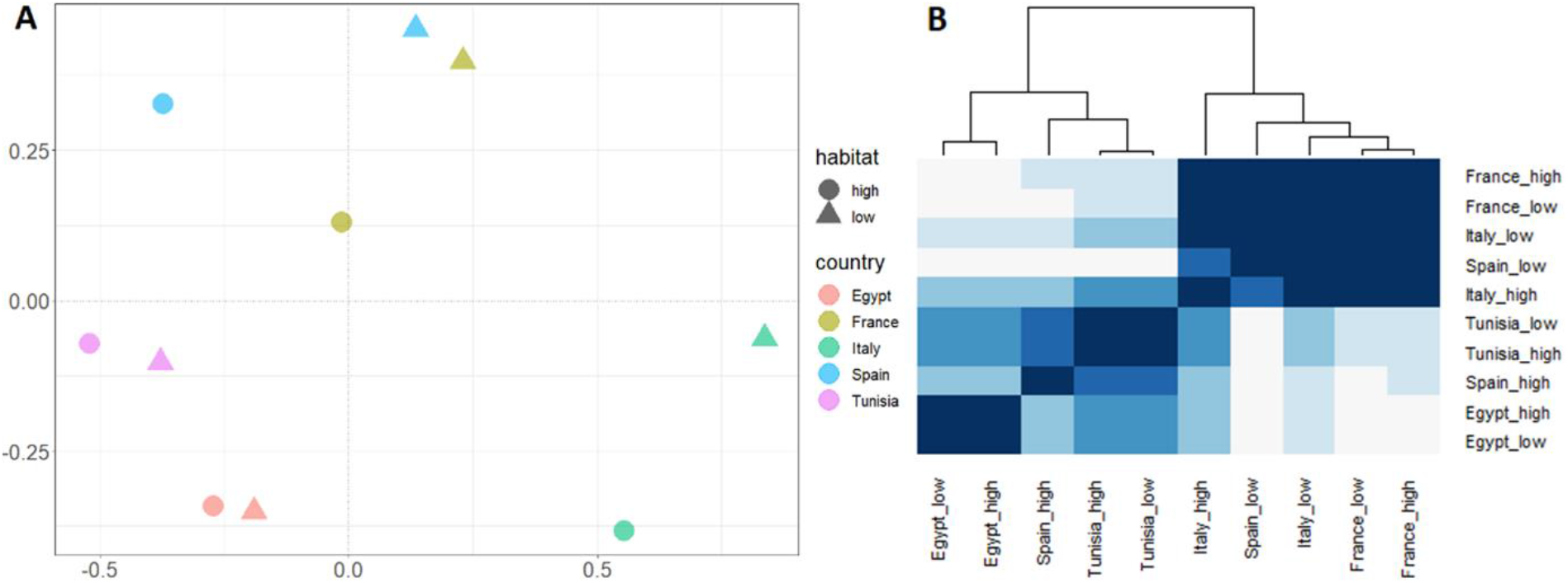
Influence of country and anthropization level (habitat) on virome structure. (A) Non-metric multidimensional scaling plot on Bray-Curtis dissimilarities of the viromes in the ten libraries. (B) Heatmap showing Pearson correlation coefficients between the viromes in the ten libraries. A dendrogram based on hierarchical clustering is shown on the top. Tile color represents Pearson coefficients, with color becoming darker along with the Pearson coefficient.

### Virus phylogeny does not suggest strict isolation of viromes from different continents

The observed differences in virome assembly could be explained, among others, by a limited connectivity between the viromes of *C. pipiens* in Africa and Europe. We explored this question using a phylogenetic approach. The existence of the Mediterranean Sea for over five million years should have allowed for isolation, and thus divergent evolution, of virus populations located in Africa and Europe. Sequences of three taxonomically-distant viruses (Culex Bunyavirus 2, Alphamesonivirus 1 and Culex Iflavi-like virus 4) from each country, together with sequences from other countries in the GenBank database, were used to generate phylogenetic trees (Fig. 8). Sequence clustering based on geography was only clear for Culex Bunyavirus 2.

**Figure 8.**
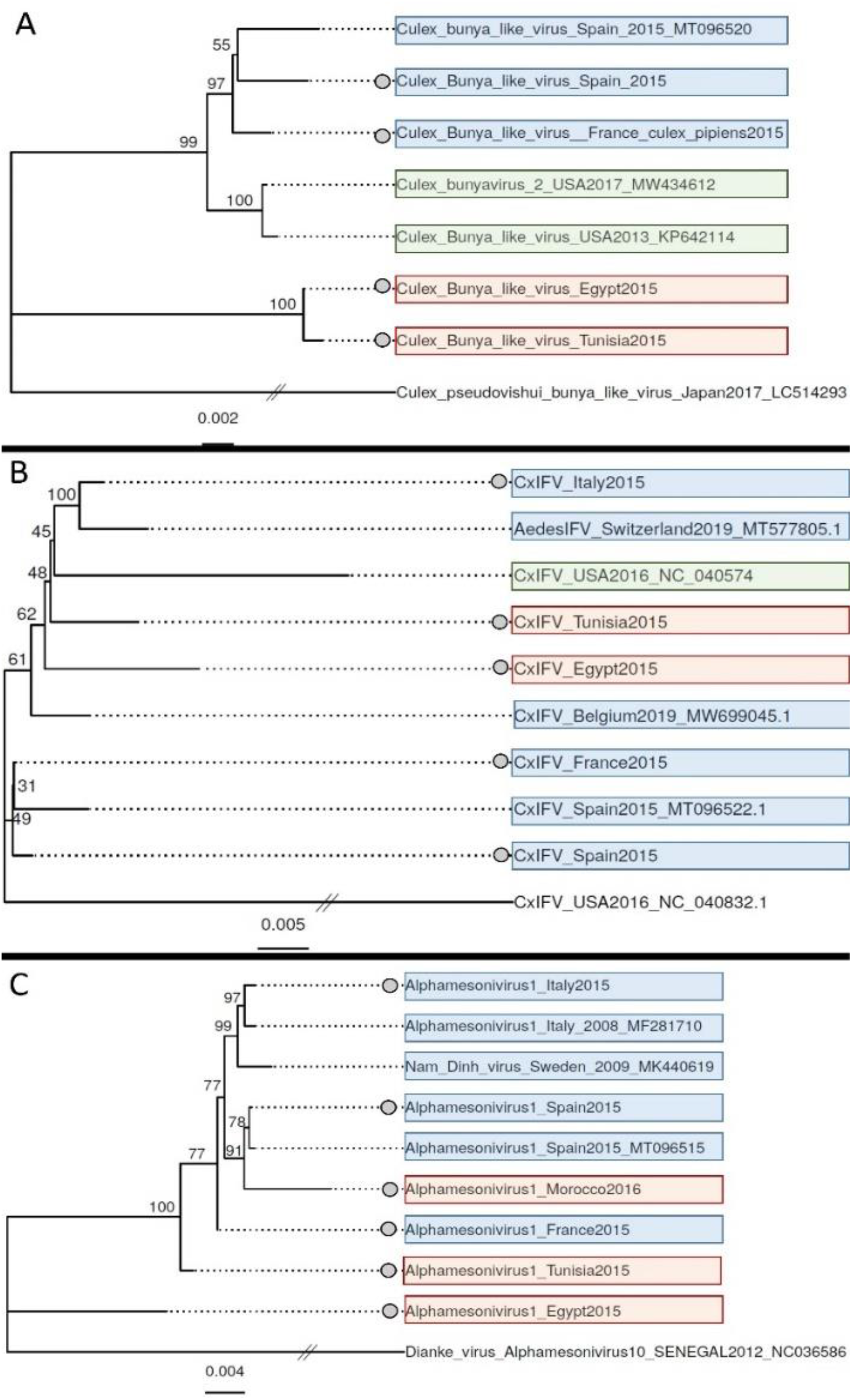
Maximum-likelihood phylogenetic trees of three viral operational taxonomic units: Culex.Bunyavirus.2__98 (A), Culex.Iflavi_like.virus.4__96 (CxIFV) (B), and Alphamesonivirus.1__97 (C). Sequences from this study are marked with a grey dot on the left of the sequence name. A divergent species from the same family was used as outgroup to root each tree (lowest branch in all the trees). The scale bar represents the number of nucleotide substitutions per site. Background color stands for geographical origin (Europe: blue, Africa: red, America: green). GenBank accession numbers of sequences from other studies are provided on the branch names.

## Discussion

Our metagenomics analysis showed a relatively high taxon conservation in the eukaryotic virome of *C. pipiens* over a large geographical area. Conserved taxa were phylogenetically distant and represented a large fraction of the virome in terms of read abundancy. Moreover, taxon conservation beyond the study region was supported by a literature analysis. This degree of taxon conservation is striking given the experimental design. Here, sampling involved mosquito populations separated by an important physical barrier, the Mediterranean Sea. Furthermore, the populations from each country were analyzed separately according to habitat anthropization, a factor that impacts virus diversity (59). This sampling design should favor differences in diversity between samples.

To our knowledge, only one study has performed an in-depth comparison of the virome of a mosquito species over a large geographical scale (28). Shi and colleagues compared the viromes from field populations of *Aedes aegypti* from North America, Africa, Asia and Oceania. They also observed numerous and diverse taxa conserved over several continents. Interestingly, the core virome in *A. aegypti* does not fully overlap that observed in *C. pipiens* here (Fig. S6). This observation represents the first evidence of specificities in the core virome of different mosquito species when compared over large geographical ranges. Hence, our results suggest not only that a core virome may be a common feature of the Culicidae, but also that core viromes could differ among mosquito species.

Two putative processes, non-mutually exclusive, could explain the observed taxon conservation in the virome of *C. pipiens*. First, the conserved virus taxa may have spread along with *C. pipiens* during geographical range expansions of the latter (40). However, the observed low similarity in the virome among *C. pipiens* individuals should hamper virome conservation during migration. Such conservation would have required the participation in migrations of a large number of *C. pipiens* individuals (each with different viruses), or specific individuals carrying all core viruses. The second process allowing taxon conservation involves the spread of the highly-conserved viruses by host species other than *C. pipiens*. For example, several mosquito species have colonized parts of the *C. pipiens* distribution (60). This second process requires that conserved viruses can be horizontally transmitted between mosquito species, a transmission mode that is rarely invoked or demonstrated in mosquito-specific viruses. In line with the second process, Alphamesonivirus 1, the best studied of the highly-conserved viruses, has a host range including mosquito species beyond the *Culex* genus and a large geographical range, although its transmission routes remain elusive (61). Currently, we cannot disentangle the relative weight of the proposed processes in taxon conservation in the *C. pipiens* virome. This situation is due to the paucity of data on the host range, transmission modes and evolutionary history for most of the highly-conserved viruses (19).

Contrary to the observations at the between-country scale, virome conservation was limited at the within-country scale. Our results thus suggest a new phenomenon: the level of taxon conservation in a mosquito virome can depend on the spatial scale, with conservation decreasing along with scale. At this point, we cannot provide a simple mechanistic explanation for this phenomenon. Whether this phenomenon takes place in the eukaryotic virome of other mosquitoes or terrestrial animals is yet unclear due to the limited number of studies robustly exploring different scales (6, 8, 62). A previous study on the mosquito virome, geographically restricted to a Caribbean island, suggests that other situations are possible (27). Shi and colleagues showed infection rates above 90% for three viruses screened in 72 *A. aegypti* adults, and similar results were found in 24 *Culex quinquefasciatus* adults.

The co-infection rates in individual mosquitoes were analyzed to explore potential non-random co-infection of certain viruses. Contrary to previous studies (27, 37, 63), a large number of individuals was tested, thus improving the analysis robustness. Mosquitoes were often infected by more than one virus, as previously observed in *C. pipiens* and other mosquito species (27, 37, 63). However, co-infection by most virus couples did not deviate from random co-occurrence. Thus, the highly-conserved viruses included in our analysis do not seem to be always simultaneously transmitted. This observation is not expected in a core virome dominated by vertically-transmitted viruses, a widespread view of the mosquito virome (19, 34).

Our prevalence data from individual mosquitoes also provided insights into an open question in virome ecology, that of the potential similarity in assembly patterns between the virome and the bacteriome (35, 64). Here, the low prevalence of certain highly-conserved viruses contrasts the high prevalence of core taxa in the bacteriome of mosquitoes, including *C. pipiens* (65, 66).

Our analyses detected an influence of geography on virome assembly in *C. pipiens*, with viromes clustering depending on their African or European origin. Jaccard and Bray-Curtis indexes suggested that clustering due to geography was mainly due to differences in relative abundancy between vOTUs, although specificities in taxonomic diversity were also observed. Moreover, the virus phylogenies obtained did not suggest a strict isolation of virus populations in the study area (Fig. 8). Our results thus suggest that viruses are shared and exchanged in the study area but, that, the resulting viromes tend to differ in read abundancy due to unknown factors.

The observed patterns in virome assembly require further support due to limits in our experimental design. First, our list of highly-conserved taxa may not be exhaustive. Our meta-transcriptomics approach is considered to be among the most sensitive for virus detection (67), and the number of mosquitoes was higher than in other studies (10, 27). Nevertheless, we cannot exclude that highly-conserved viruses have not been detected due to, among others, low viral loads or temporal dynamics of certain viruses. Secondly, certain highly-conserved taxa may not be fully conserved if the spatial or temporal extent of the sampling is increased. For example, the number of libraries number used in our metagenomics analysis is in the same range as those found in most studies on the mosquito virome (10, 34-36, 38) but yet limited. Furthermore, our study only analyzed adult females. Whether assembly patterns in the mosquito virome are conserved along the life cycle or between sexes are questions that remain poorly studied (28, 34). Thirdly, the analyses of infection rates involved a limited set of viruses. We chose to include the two highly-conserved viruses with read counts and contig lengths warranting a robust detection. However, other highly-conserved viruses may always have high infection rates in *C. pipiens* populations. Future work on a larger panel of highly-conserved viruses could thus unveil the coexistence of contrasted infection rates. Another point that requires further study is the potential influence of habitat anthropization on the mosquito virome. Here, the limited number of libraries did not allow for a robust analysis.

Uncovering deterministic patterns in the eukaryotic viromes of terrestrial animals is considered to be challenging (6, 62). This study shows that integrating different spatial scales allows to identify deterministic patterns in the *C. pipiens* virome. Our data provide predictive power and can represent a base for hypothesis-driven studies on, among others, the potential role of the *C. pipiens* virome in pathogen transmission. For example, the highly-conserved viruses identified here represent candidates for studies on their influence on pathogen transmission. Moreover, the virome differences observed among individuals or regions could participate to the complex epidemiological patterns associated to mosquito-borne arboviruses.

## Material and Methods

### Mosquito sampling

Mosquitoes were captured in five countries (Egypt, France, Italy, Spain and Tunisia; Fig. 1). The genetics of the *C. pipiens* complex have been shown to be relatively homogeneous in the region (42). In each country, mosquitoes were sampled in several sites (Fig. S7 and Table S4). Sites were representative of habitats of *C. pipiens* in the study region, including marshlands, riverine woodlands, crop fields or urbanized areas. Sites were grouped into two categories based on habitat anthropization (Table S4). The two categories were broadly defined to take into account differences in the diversity and availability of habitats between countries. A “high” anthropization level was broadly defined as a peri-urban site, located less than 500 m from the perimeter of a densely populated area. Similarly, a “low” level was largely defined as any agricultural or natural site with a low population density.

Several mosquito collections were carried out in each site between April and October in 2015. After overnight trapping, mainly using CO2-baited CDC-type traps, mosquitoes were identified to the species level on a chill table using morphological determination keys. Non-engorged females of *C. pipiens sensu lato* were distributed in pools of around 30 individuals (Table S5), and stored at -80 °C until further processing. An additional sampling was carried out in France in 2018. The latter sampling followed the same protocol as in 2015 but for pool size (pools of six females or individual females, Table S5).

### Library preparation and sequencing

Samples were analyzed using meta-transcriptomics (12). Ten libraries (one library per country and anthropization level) were prepared for Illumina sequencing as previously described (43). Briefly, mosquito pools were homogenized in ice-cold 1X PBS (Tissue Lyser, Qiagen). An aliquot of 150 µl was used for total RNA isolation with the Nucleospin RNA virus kit (Macherey Nagel). Ten RNA samples were generated by pooling aliquots of RNA extractions from all the pools from the same country and anthropization level. The ten samples were retro-transcribed to cDNA (SuperScript IV reverse transcriptase, Invitrogen) with random hexamers. Then, cDNA was amplified using multiple displacement amplification with phi29 polymerase and random hexamers. Libraries were prepared with the TruSeq DNA Nano Library Prep kit (Illumina). Libraries were sequenced at an expected depth of 80 million reads on a HiSeq2000 (Illumina) in a 150-bp single-read format. Library preparations and sequencing were outsourced to DNAVision (Charleroi, Belgium).

### Generation of viral operational taxonomic units

We used an updated version of the SnakeVir bioinformatics pipeline to identify potential viral sequences from reads (44). Briefly, library adapters and low-quality sequences in reads were removed with Cutadapt 1.6 (Martin, 2011). Reads from rRNA sequences from diptera and bacteria were then removed through mapping with BWA 0.7.15 (45) against SILVA rRNA bases. The remaining reads were *de-novo* assembled with Megahit version 1.1.2 (46). The resulting contigs were used in a second *de-novo* assembly with CAP3 (Huang, 1999). A contig was considered a virus-like sequence if a viral sequence was found as the first best hit by Diamond (Buchfink et al., 2015) after comparison with the NCBI nr database (release 234, 10/2019; e-value cutoff = 10^−3^). Then, virus-like contigs were screened for sequences potentially derived from endogenous viral elements (EVEs) using BLASTn against NCBI nt (release 249, 6/2022; e-value cutoff = 10^−3^). Contigs with a non-viral best hit with a coverage above 25% were considered as potential EVEs and removed.

The number of reads per virus-like contig and library was quantified through mapping with BWA 0.7.15 (45). Duplicate reads had been previously removed with the markdup tool in Samtools (47). Contigs sharing the same best hit were grouped into a viral operational taxonomic unit (vOTU) as previously described (27, 44). Names of vOTUs were generated through combining the best-hit name and the mean percent identity at the amino-acid level of the contigs with the best hit. The percent identity was placed at the end of the best-hit name after a double underscore. The vOTUs with percent identities above 90 % can be considered to include the best-hit species. Spaces and hyphens in vOTU names were replaced by dots and underscores respectively to facilitate analysis in R software.

The taxonomy of vOTUs was generated with SnakeVir (44). To avoid incomplete virus taxonomies, the vOTUs were grouped into an arbitrary taxonomic level, here called “cluster”, similar to the Family rank (44). The cluster was defined as the family of the best hit whenever available in GenBank databases. For those best hits without a full taxonomic classification, the most likely family was defined from the associated literature and BLAST searches. Clusters were named using the family name without the “viridae” suffix. If analyses failed to indicate a likely family for a given vOTU, the cluster was generated as the name of the most probable Order of the vOTU and replacing the “virales” suffix with “_unclass”.

### Validation of viral operational taxonomic units

Two arbitrary detection thresholds were applied to the list of vOTUs as previously described (44). First, a vOTUs was considered non-detected in a given library when the number of reads was below ten. Consequently, the read number was set to zero in the library. After this step, vOTUs with less than 100 reads in total were removed. We next validated that taxon conservation was not due to contamination during sample processing and sequencing. To do so, the vOTUs in the ten libraries of this study were compared to those found in six libraries sequenced in parallel and processed in the same laboratory for another project. These libraries derived from adult females of the mosquito *Aedes vexans* from Senegal and have been previously described (44). This comparison found 25 vOTUs in common. Twelve vOTUs were not considered as potential contaminants because they had a mean read abundance at least two logs higher in the *C. pipiens* libraries. None of the remaining vOTUs was part of the most prevalent vOTUs in the *C. pipiens* libraries (Table S2), or was present in all libraries.

### Analysis of published viromes of *Culex pipiens*

A search for articles with robust characterizations of the virome of *C. pipiens* was done in PubMed (October, 2021). The criteria defined to select datasets included: (i) the minimum number of mosquito individuals was set to 100, (ii) the minimal sequencing depth was set to 1 million reads per library, and (iii) the virome diversity should be dominated by viruses from arthropods. Four studies were selected for the analysis (10, 38, 48, 49). All studies analyzed populations in a single country (China, Morocco, Serbia or Sweden; Table S3). There were important methodological differences among studies (Table S3). For example, the studies widely differed in mosquito number, library preparation or sequencing depth. Moreover, the studies had mosquito numbers, sequencing depths or a yield in virus-like reads largely below the ones in this study. Hence, we limited the comparison to the presence of the highly-conserved taxons found in our dataset.

### PCR detection of selected virus taxa

We developed RT-qPCRs using SYBR green chemistry for the detection of several vOTUs. First, we selected vOTUs in four highly-conserved clusters - the Mesoni, Ifla, Nege and Reo clusters – for the analysis. We also selected vOTUs in two additional clusters present in nine libraries, the Chu and Bunya_unclass clusters. Primer design was done with LightCycler Probe design Software 2.0 (Roche) and based on the contigs of each vOTU. Up to three primer pairs were tested per vOTU. The PCR output was not satisfactory in terms of specificity and sensibility for vOTUs in the Reo (not shown), and vOTUs of this cluster were not included in the analysis. Moreover, the PCR design for the most prevalent vOTU in the Nege was not satisfactory (Dezidougou.virus__98; not shown) and was discarded from the analysis. A satisfactory PCR design was obtained for Culex.negev_like.virus.1__99, the second most prevalent vOTU in the Nege. Culex.negev_like.virus.1__99 was thus included in the analysis. The final set of five vOTUs and their prevalence among libraries are presented in Figure S2. The primer pairs can be found in Table S6.

RT-qPCR conditions were as follows. First, total RNA was reverse transcribed using the RevertAid First Strain cDNA kit (ThermoFisher) and random primers following manufacturer’s instructions. Then, qPCR amplifications were carried out with the Brilliant III-Sybrgreen QPCR Master Mix (Agilent) in a total volume of 20 µl consisting of 2 µl of cDNA template, 0.3 µM of the forward and reverse primers, 10 µl of Master Mix 2X and RNase-free H_2_O. Thermal cycling was 95°C for 10 min, 40 cycles of 95°C for 20 s, 60°C for 20 s (55° for Culex.Bunyavirus.2__96), and a melting curve analysis involving 95°C for 10 s, 65°C for 60 s and 97°C for 1 s. Amplicons were Sanger sequenced (Genewiz, Germany) whenever an unclear melting curve was observed.

### Phylogenetic analysis

First, vOTUs were selected for phylogenetic analysis only if detected in at least two European countries and the two African countries. Moreover, for a given vOTU, contigs from the different countries had to encompass the same genomic region with at least 10X read coverage over 500 bp. This analysis selected three vOTUs: Alphamesonivirus.1__97, Culex.Iflavi_like.virus.4__96 and Culex.Bunyavirus.2__96. The contigs of those vOTUs had mean identities at the nucleotide level above 90% with their best hits. The contigs in the vOTUs were thus considered as belonging to the same species as their best hits.

The contigs selected for Alphamesonivirus.1__97 and Culex.Iflavi_like.virus.4__96 represented nearly-full genomes. The contigs of Culex.Bunyavirus.2__96 represented a nearly-complete L segment. In addition to those contigs, homologous sequences from different countries found in GenBank were also included. The search for homologous sequences unveiled that sequences from Culex picorna-like virus 1 and Aedes Ifla-like virus belong to the same species as Culex Iflavi-like virus 4. Similarly, sequences from Culex Bunya-like virus and Bunyanwera environmental sample belong to the same species as Culex Bunyavirus 2.

Alignments were generated with Mega X using the ClustalW algorithm for each group of sequences (50). All ambiguously aligned regions were removed using TrimAL (51). The alignments had 16000, 8800 and 7000 nucleotide positions for the sequences from Alphamesonivirus 1, Culex Iflavi-like virus 4 and Culex Bunyavirus 2 respectively. For each alignment, IQ-Tree (v.1.6.1) (52) and its module *modelfinder* was used to determine the best-fit model of nucleotide substitution. The TN+F+R3 model (Tamura and Nei substitution model with empirical base frequency and Gamma distributions for three categories) was selected for each alignment. Maximum likelihood phylogenetic trees were then produced with 1000 bootstrapping replications. Trees were visualized and annotated with the R package *ggtree* (53).

### Statistical analyses

Statistical analyses are described along the text and were conducted using R version 3.5.1. The associated figures were generated with the *ggplot2* package. Statistical analyses on virome data were done, unless specifically stated, at the cluster level to avoid potential biases in vOTU identification. The Jaccard and Bray-Curtis dissimilarities were estimated using the *vegan* package (54) from data normalized using Total Sum Normalization (55). Rarefaction curves were generated with the *vegan* package (54). Maximum-likelihood estimates of infection rates were obtained with the *binGroup* package (56). A probabilistic analysis of virus co-occurrence was done with the *coocur* package (57).

### Data availability

Raw reads and selected contigs were deposited in GenBank associated to BioProject PRJNA806751.

## Acknowledgements

We thank the personnel of the EID Méditerranée (agencies of Arles, Canet-en-Roussillon, Frejorgues and Montcalm) and Lou Carlier for their involvement in mosquito collection. We also thank Emmanuel Albina for his contribution in the initial phases of the project. This study was partially funded by EU grant FP7-613996 Vmerge and a grant from the Montpellier University of Excellence program (MUSE, ArboSud project). Moreover, SG was recipient of funding from the Direction Générale de l’Alimentation from the French Ministry of Agriculture and Food (DGAl grant agreement: SPA17 n◦0079-E). The contents of this publication are the sole responsibility of the authors and do not necessarily reflect the views of the European Commission.

