## Supplementary information for "Spatial scale influences taxon conservation in the eukaryotic virome of a mosquito"

Table S6 .......... 7
Supplementary Annex 1. Gsub tool for sequence submission to GenBank .......... 9

Figure S1. Rarefaction curves for viral operational taxonomic units (vOTUs) of each library. No filtering step was applied to the vOTU dataset. Abscise axis is limited to $4x10^6$ reads to improve curve visualization.

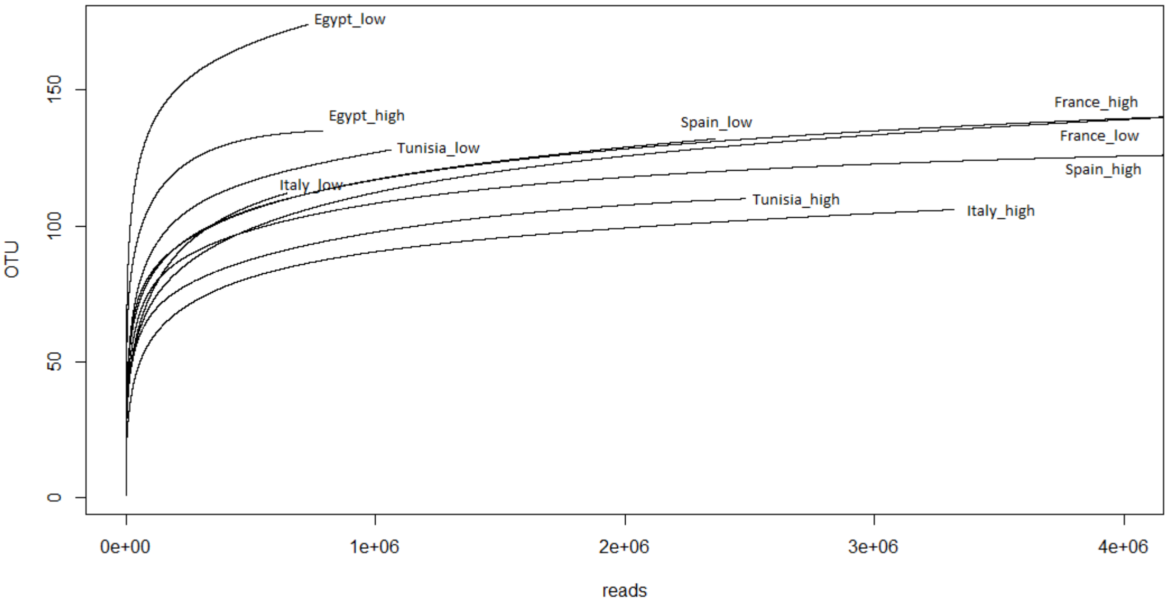

Figure S2. Distribution of the mean percent identity at the amino acid level of viral operational taxonomic units and their best hit. Most of the percent identity values felt below 90%, strongly suggesting a predominance of new virus species in our dataset.

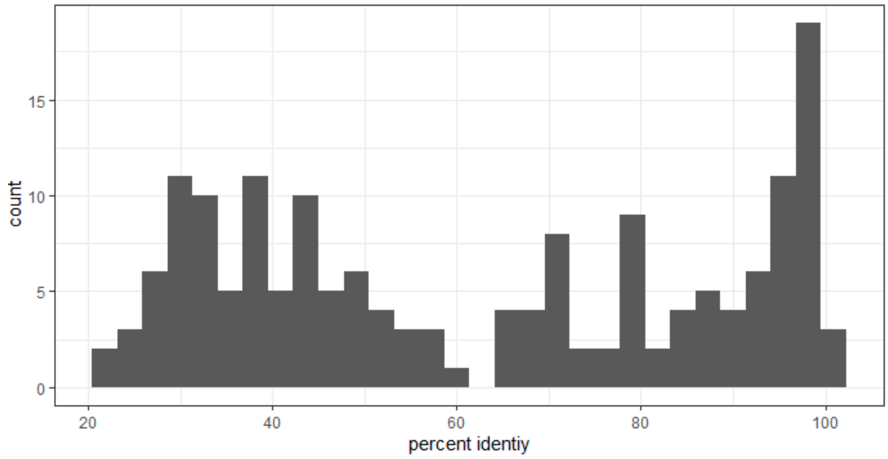

Figure S3. Prevalence in the metagenomics libraries of the five viral operational taxonomic units (vOTUs) selected for PCR screening. The vOTU names appear on the left and their cluster on the rigth. Coloured tiles indicate detection in a given library (y-axis). Colour in tiles is specific to each vOTU to facilitated comparison.

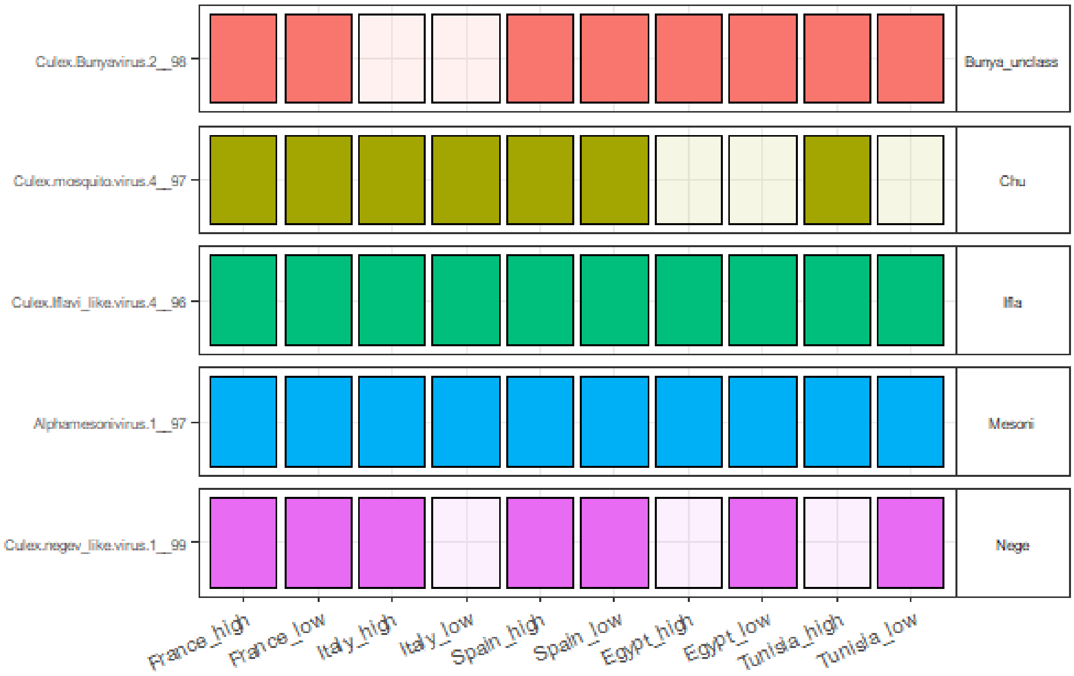

Figure S4. Taxon richness, Simpson's and Shannon's indexes for each library presented depending on (A) Continent (Europe: N, Africa: S), (B) anthropization level (high and low), (C) mosquito number per library and (D) sampling sites per library. All analyses were done at the cluster level. None of the four variables had a significant effect on the three indexes. Dot color stands for country (Egypt: red, France: light brown, Italy: green, Spain: blue, Tunisia: magenta). Dot shape stands for anthropization level (high: circle, low: triangle).

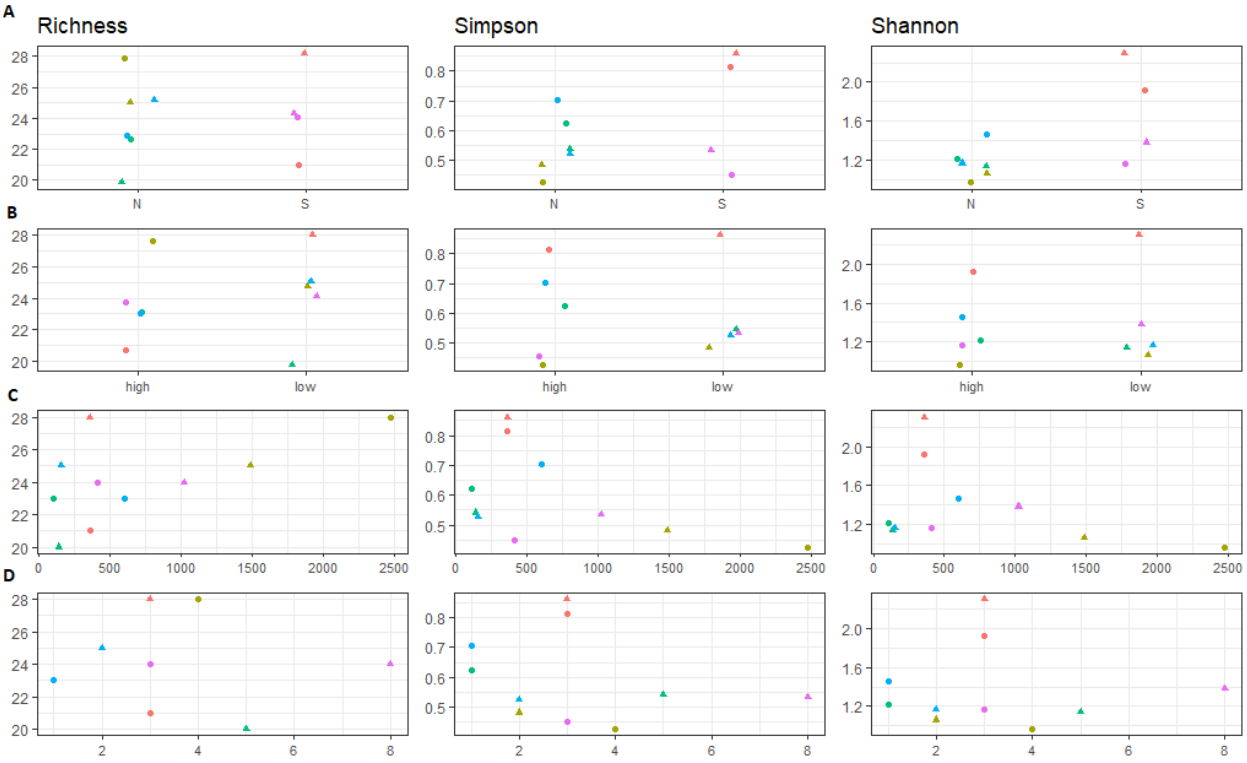

Figure S5. (A) Bray-Curtis and Jaccard dissimilarities among the viromes found in the ten libraries. (B) and (C): the same dissimilarities are shown following region or anthropization level, respectively. Colors in the “same” level indicate region (Europe: blue, Africa: red) or anthropization level (high: blue, low: red) in the B and C panels respectively. Bray-Curtis dissimilarities were significantly lower whenever viromes were from the same continent (Region) but Jaccard dissimilarities were not (Kruskall Wallis test, p = 10E-3 and 0.25, respectively; Fig. S7B). No significant difference was found for any of the two dissimilarities depending on anthropization level of the habitat (Kruskall Wallis test, p > 0.61; Fig. S7C).

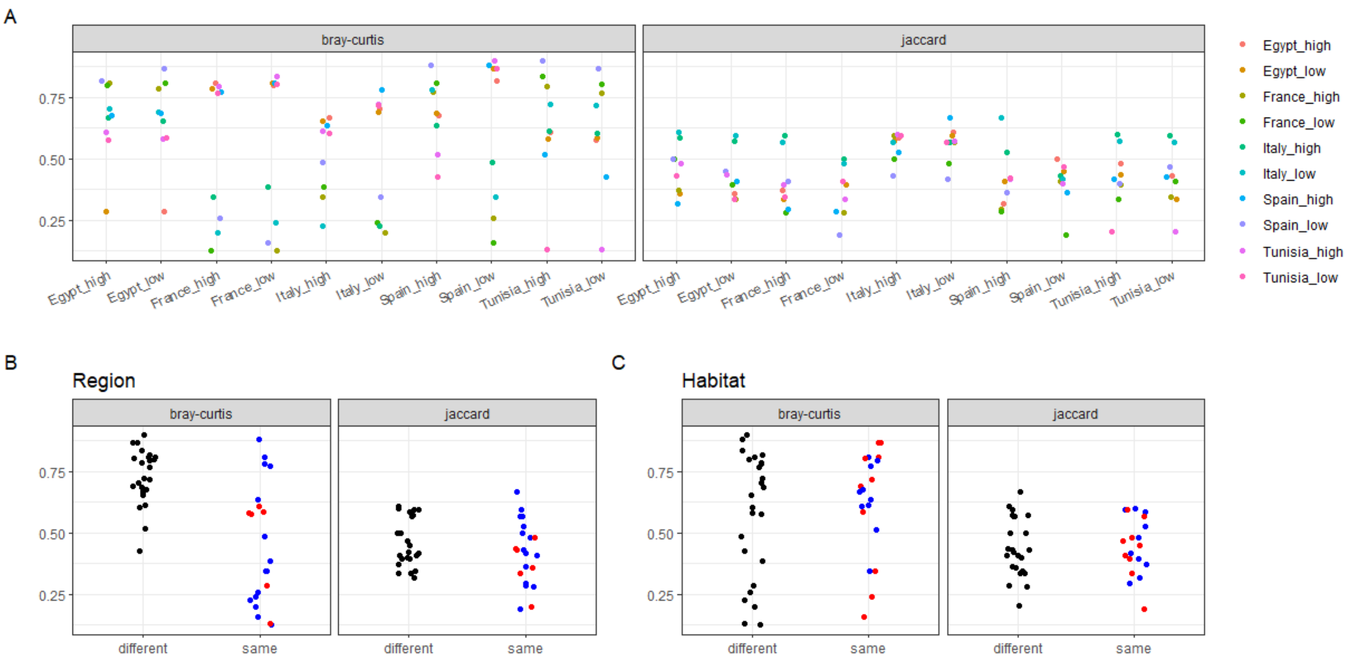

Figure S6. Heatmap with the conserved viral clusters in the viromes of *Culex pipiens* and *Aedes aegypti*. Clusters present in all countries in our dataset (n = 18) were compared to those found associated to *A. aegypti* in at least two continents (n = 12; data from Shi et al. 2020. *Microorganisms*). Grey tiles indicate conservation in a single mosquito species and black tiles indicate conservation in the two species. Since the cluster rank is arbitrary, the Order rank of each cluster is provided in the last column.

| **Cluster** | ***C. pipiens*** | ***A. aegypti*** | **Order** |
|---|---|---|---|
| Bunya_unclass | | | Bunyavirales |
| Chryso | | | Ghabrivirales |
| Chu | | | Jingchuvirales |
| Flavi | | | Amarillovirales |
| Ifla | | | Picornavirales |
| Luteo_unclass | | | Luteo-Sobemo_order |
| Martelli_unclass | | | Martellivirales |
| Mesoni | | | Nidovirales |
| Meta | | | Ortevirales |
| Mononega_unclass | | | Mononegavirales |
| Nege | | | Martellivirales |
| Noda | | | Nodamuvirales |
| Orthomyxo | | | Articulavirales |
| Partiti | | | Durnavirales |
| Permutotetra | | | Permutotetra_order |
| Phasma | | | Bunyavirales |
| Phenui | | | Bunyavirales |
| Picorna_unclass | | | Picornavirales |
| Reo | | | Reovirales |
| Rhabdo | | | Mononegavirales |
| Toti | | | Ghabrivirales |
| Virga | | | Martellivirales |
| Xinmo | | | Mononegavirales |

Figure S7. Mosquito sampling sites in each country: (A) Egypt, (B) Spain, (C) Tunisia, (D) France, (E) Italy.

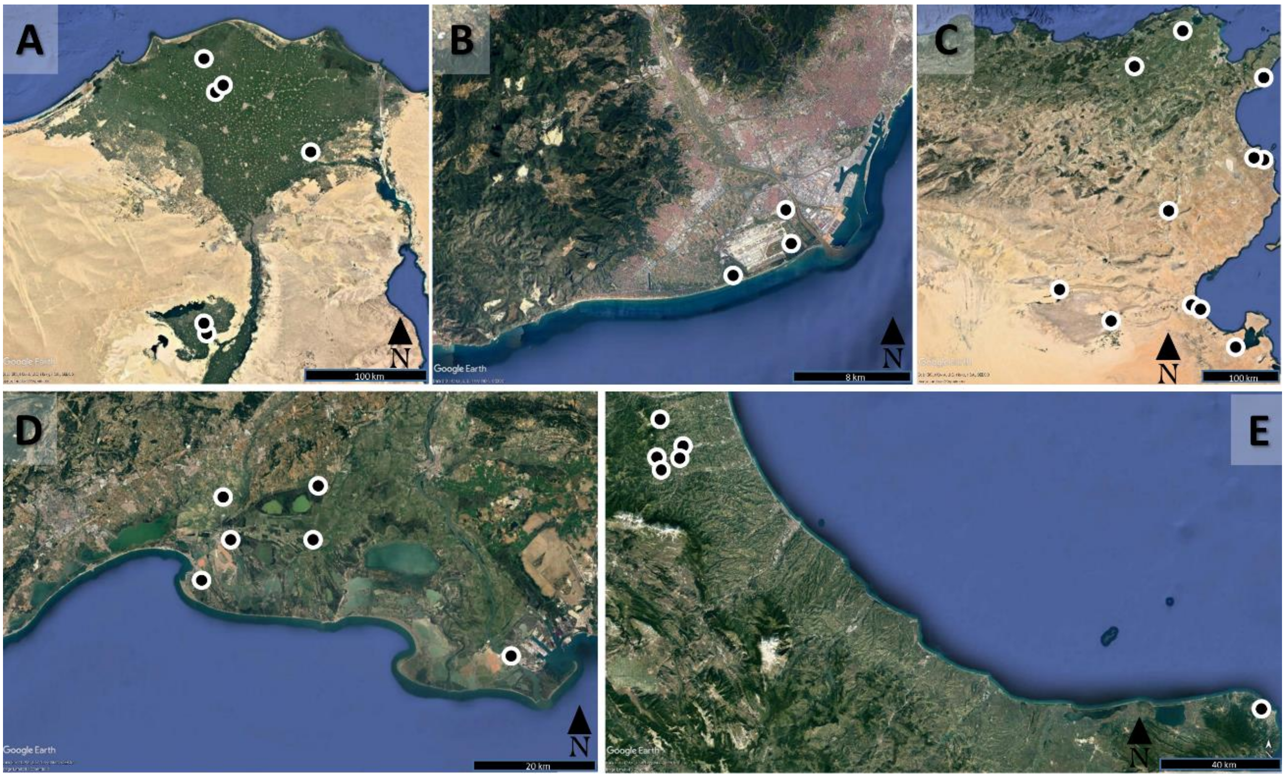

Table S1. Metadata and sequencing output for the ten libraries of this study.

| library | Country | Anthropisation | Sites | Mosquitoes | Reads x $10^7$ | Viral reads x $10^5$ |
|---|---|---|---|---|---|---|
| Egypt_high | Egypt | high | 3 | 360 | 6.24 | 7.88 |
| Egypt_low | Egypt | low | 3 | 360 | 6.63 | 7.27 |
| France_high | France | high | 4 | 2475 | 6.97 | 97.70 |
| France_low | France | low | 2 | 1489 | 8.03 | 143.37 |
| Italy_high | Italy | high | 1 | 105 | 8.26 | 33.19 |
| Italy_low | Italy | low | 5 | 137 | 7.73 | 6.44 |
| Spain_high | Spain | high | 1 | 600 | 7.56 | 59.56 |
| Spain_low | Spain | low | 2 | 155 | 7.99 | 23.60 |
| Tunisia_high | Tunisia | high | 3 | 411 | 6.57 | 24.84 |
| Tunisia_low | Tunisia | low | 8 | 1023 | 5.54 | 10.63 |

Table S2. Viral operational taxonomic units (vOTUs) with the largest prevalence in each highly-conserved cluster. The libs_OTU column provides the number of libraries with a positive detection.

| **vOTU** | **cluster** | **libs_OTU** |
|---|---|---|
| Alphamesonivirus.1__97 | Mesoni | 10 |
| Culex.Iflavi_like.virus.4__96 | Ifla | 10 |
| Culex.mononega_like.virus.2__97 | Mononega_unclass | 8 |
| Culex.mosquito.virus.1__99 | Noda | 10 |
| Culex_associated.Luteo_like.virus__86 | Luteo_unclass | 10 |
| Dezidougou.virus__98 | Nege | 8 |
| Hubei.diptera.virus.20__25 | Reo | 10 |
| Hubei.tetragnatha.maxillosa.virus.8__43 | Partiti | 9 |
| Hubei.virga_like.virus.2__79 | Martelli_unclass | 10 |
| Nephila.clavipes.virus.2__45 | Picorna_unclass | 10 |

Table S3. Main parameters of the studies used in the virome comparison. Whenever several mosquito species had been analyzed in a study, only the libraries from *Cx. pipiens* individuals were used to estimate parameter values. The values of this study are provided for comparison. Raw and virus-like reads are presented in millions. NP: not provided.

| **country** | **mosquitoes** | **libraries** | **raw reads** | **virus-like reads** | **doi** |
|---|---|---|---|---|---|
| China | 10497 | 6 | 167.7 | 1.6 | 10.1371/journal.pntd.0009381 |
| Morocco | 1488 | 3 | 11.4 | 2.4 | 10.1016/j.virol.2019.02.007 |
| Serbia | 360 | 1 | 43.7 | NP | 10.3390/v12090975 |
| Sweden | 120 | 6 | 253.8 | 1.4 | 10.3390/v11111033 |
| This study | 7115 | 10 | 715.2 | 41.4 | |

Table S4. Coordinates and anthropization level of the sampling sites.

| **country** | **site** | **latitude** | **longitude** | **anthropisation level** |
|---|---|---|---|---|
| Egypt | Kafr Elshiekh-hospital | 31° 06' 59.7"N | 30° 56' 33.0"E | high |
| Egypt | Al Fayoum-Water station | 29° 18' 30.1"N | 30° 50' 34.2"E | high |
| Egypt | Al Qorin | 30° 36' 48.4"N | 31° 44' 30.9"E | high |
| Egypt | Qarajah | 31° 07' 22.9"N | 30° 59' 04.3"E | low |

| | | | | |
|---|---|---|---|---|
| Egypt | Ezbet El-Sawy | 31° 17' 25.6"N | 30° 49' 16.3"E | low |
| Egypt | Monshaat Abd Allah | 29° 20' 01.8"N | 30° 50' 26.2"E | low |
| France | Bois François | 43° 23' 56.39"N | 4° 47' 10.20"E | high |
| France | Jasse brulée | 43° 37' 58.24"N | 4° 12' 22.59"E | high |
| France | L'Elysette | 43° 30' 57.87"N | 4° 9' 20.48"E | high |
| France | Mas d'Avon | 43° 34' 5.14"N | 4° 12' 29.24"E | high |
| France | Petite Sylve | 43° 34' 16.56"N | 4° 22' 56.07"E | low |
| France | Bois d'Espeyran | 43° 38' 56.88"N | 4° 23' 36.91"E | low |
| Italy | Teramo | 42° 39' 16.19"N | 13° 42' 38.21"E | high |
| Italy | Gattia | 42° 38' 17.00"N | 13° 40' 5.69"E | low |
| Italy | Piano grande | 42° 39' 1.07"N | 13° 38' 58.00"E | low |
| Italy | Colleatterrato Alto | 42° 41' 25.89"N | 13° 44' 28.28"E | low |
| Italy | Villa Lempa | 42° 47' 1.97"N | 13° 39' 5.25"E | low |
| Italy | Vieste | 41° 54' 42.84"N | 16° 7' 21.86"E | low |
| Spain | Prat de Llobregat | 41° 18' 59.76"N | 2° 6' 15.35"E | high |
| Spain | La Ricarda | 41° 18' 2.27"N | 2° 6' 22.11"E | low |
| Spain | El Remolar | 41° 16' 48.10"N | 2° 3' 51.51"E | low |
| Tunisia | GABES_1 | 33° 53' 49.2000'' N | 10° 5' 49.2000'' E | high |
| Tunisia | MOKNINE_1 | 35° 37' 46.5734'' N | 10° 52' 58.3398'' E | high |
| Tunisia | TAZARKA | 36° 31' 10.2842'' N | 10° 49' 57.8649'' E | high |
| Tunisia | MATEUR | 37° 03.790' N | 9° 39.641' E | low |
| Tunisia | SBZ | 35° 0' 18.0000'' N | 9° 37' 22.8000'' E | low |
| Tunisia | MEDENINE | 33° 29' 54.4840'' N | 10° 38' 17.2169'' E | low |
| Tunisia | KEBILI | 33° 44' 12.3880'' N | 8° 53' 53.2819'' E | low |
| Tunisia | GABES_2 | 33° 54' 40.8016'' N | 10° 3' 9.9722'' E | low |
| Tunisia | BOUSSALEM | 36° 38' 26.8548'' N | 9° 0' 42.1452'' E | low |
| Tunisia | TOZEUR | 33° 59.660' N | 8° 09.630' E | low |
| Tunisia | MOKNINE_2 | 35° 37' 22.8'' N | 10° 53' 20.4'' E | low |

Table S5. Sample sizes used in the analysis of infection rates. The column "mosquitoes" provides number of adult females of *Culex pipiens*. The batch of pools from France in 2018 included 184 pools of a single female that were used in the co-infection analysis. The mosquito samples from Morocco derived from another study (Bennouna et al., 2019, *Virology*).

| country | year | mosquitoes | pools | pool size |
|---|---|---|---|---|
| Egypt | 2015 | 630 | 21 | 30 |
| France | 2015 | 3764 | 131 | 6 to 34 |
| France | 2018 | 748 | 278 | 1 to 6 |
| Italy | 2015 | 142 | 16 | 4 to 30 |
| Morocco | 2016 | 1349 | 44 | 20 to 35 |
| Spain | 2015 | 780 | 26 | 30 |
| Tunisia | 2015 | 577 | 27 | 4 to 30 |

Table S6. Primer pairs used in vOTU detection in mosquitoes with RT-qPCR. Sequences are provided in the 5'-3' direction. The target region in the respective GenBank accession is provided.

| Cluster | vOTU | primer | sequence | Target region | Accession |
|---|---|---|---|---|---|
| Mesoni | Alphamesonivirus.1__97 | Meso_F469 | GCCTACACTGTACATAACATACG | 463-485 | KU095838 |
| Mesoni | Alphamesonivirus.1__97 | Meso_613rev | TAGATTCAATAGTGGTAAGTTCGC | 607-584 | KU095838 |
| Ifla | Culex.Iflavi_like.virus.4__96 | Picorna_fwd | GGAAAGTTTGTCTTGGT | 6439-6455 | NC040574 |
| Ifla | Culex.Iflavi_like.virus.4__96 | Picorna_rev | TTCATACCAGGAGGCC | 6647-6632 | NC040574 |
| Nege | Culex.negev_like.virus.1__99 | Nege_fwd | GTCGGAGACCCGGTTA | 6739-6754 | NC035124 |
| Nege | Culex.negev_like.virus.1__99 | Nege_rev | GTCCCAAACAGGAGCC | 6956-6936 | NC035124 |
| Chu | Culex.mosquito.virus.4__97 | Imjin_fwd | ACATGTACTCTGTCCCTA | 2781-2798 | MH188031 |
| Chu | Culex.mosquito.virus.4__97 | Imjin_rev | GTCTTGACTCTCCATTC | 3147-3131 | MH188031 |
| Bunya_unclass | Culex.Bunyavirus.2__96 | Bunya-E- Fwd | AGTGTTACTTTAAGACCCGAC | 3603-3622 | KP642114 |
| Bunya_unclass | Culex.Bunyavirus.2__96 | Bunya-E-Rev | GAGACTTGATAGCAGGCGAA | 3914-3895 | KP642114 |

### Supplementary Annex 1. Gsub tool for sequence submission to GenBank

Few tools allow easy annotation and submission of a large number of sequences from different viruses (Tisza, Belford, Dominguez-Huerta, Bolduc, & Buck, 2021). To facilitate sequence submission by non-bioinformaticiens, we developed the Gsub tool. Gsub was written in Python and uses a graphical interphase compatible with Windows operating system. All script and examples of the input files can be accessed on GitHub (https://github.com/FlorianCHA/Gsub/tree/master/Gsub).

Gsub requires three input files: (i) a FASTA file with the nucleotide sequences, (ii) a template file generated using the GenBank Submission Template tool (https://submit.ncbi.nlm.nih.gov/genbank/template/submission/), and (iii) a source file with the metadata to insert in the different fields of GenBank accession files for each sequence.

Gsub uses python package *ORFFinder* (https://github.com/Chokyotager/ORFFinder) to identify and annotate open reading frames (ORFs). Then, python package *pyHMMER* (https://pyhmmer.readthedocs.io/en/stable/index.html#) is used to detect potential polymerase-encoding ORFs (detection threshold: score > 100 and e-value > $10^{-4}$). Table2asn (https://www.ncbi.nlm.nih.gov/genbank/table2asn/) is used to collect the different pieces of information and to generate the corresponding sqn file for each sequence. The packages *Gooey* (https://github.com/chriskiehl/Gooey) and *Pyinstaller* 5.2 (https://pyinstaller.org/) allow respectively for a graphical interface and to run Gsub in Windows or Linux without installing a Python interpreter.

Gsub has an optional Phyton module, *SnakeVir2Gsub*, to facilitate the generation of the source file. SnakeVir2Gsub collects metadata and virus taxonomy directly from output files of SnakeVir. *SnakeVir2Gsub* uses a Blastn output (virus-like contigs against GenBank nt) to identify sequences of known viruses and collect their names. Sequences were defined as belonging to known viruses if bearing more than 80% identity and query cover with their Blastn best hit. The Diamond output file in SnakeVir was used to collect the genome type (DNA or RNA) and the taxonomic information (Realm and Order levels).

Gsub generates two types of files for each sequence to submit. The first file is a sqn file to be submitted to GenBank through e-mail. The sqn file contains the metadata, the annotated ORFs and their amino-acid sequences, and the nucleotide sequence. The second file is a Genbank flatfile that allows visualization in the same format as the final GenBank accession file. This second file allows to check for errors before submission. Moreover, Gsub generates a common discrepancy report for all the sequences with error messages due to discrepancies with submission requirements.
